# Genomic signatures of desert adaptation at gene-rich regions in zebu cattle from the African drylands

**DOI:** 10.1101/2021.12.15.472850

**Authors:** Abdulfatai Tijjani, Bashir Salim, Marcos Vinicius Barbosa da Silva, Hamza A. Eltahir, Taha H. Musa, Karen Marshall, Olivier Hanotte, Hassan H. Musa

## Abstract

Sudan, the largest country in Africa, acts as a corridor between North and sub-Saharan Africa along the river Niles. It comprises warm arid and semi-arid grazing lands, and it is home to the second-largest African population of indigenous livestock. Indigenous Sudanese cattle are mainly indicine/zebu (humped) type. They thrive in the harshest dryland environments characterised by high temperatures, long seasonal dry periods, nutritional shortages, and vector diseases challenges. We investigated genome diversity in six indigenous African zebu breeds sampled in Sudan (Aryashai, Baggara, Butana, Fulani, Gash, and Kenana). We adopted three genomic scan approaches to identify candidate selective sweeps regions (*ZH_p_*, *F_ST_*, XP-EHH). We identified a set of gene-rich selective sweep regions shared across African and Asian zebu or unique to Sudanese zebu. In particular, African and Asian zebu candidate gene-rich regions are detected on chromosomes 2, 5 and 7. They include genes involved in immune response, body size and conformation, and stress response to heat. In addition, a 250 kb selective sweep on chromosome 16 was detected exclusively in five Sudanese zebu populations. This region spans seven genes, including *PLCH2*, *PEX10*, *PRKCZ* and *SKI*, which are involved in alternative adaptive metabolic strategies of insulin signalling, glucose homeostasis, and fat metabolism. Together, these genes may contribute to the zebu cattle resilience to heat, nutritional and water shortages. Our results highlight the putative importance of selection at gene-rich genome regions, which might be under a common regulatory genetic control, as an evolutionary mechanism for rapid adaptation to the complexity of environmental challenges.

## Introduction

Sudan is one of the leading African countries in livestock production. The livestock sector plays a critical role in the Sudanese economy and the welfare of the whole population (Wilson, 2018). Of its 101 million livestock heads, cattle comprise approximately 41% (~41 million cattle heads). The remaining are sheep, goats and camels. According to Epstein (1971), cattle were introduced to Sudan from Asia through the Nile Valley and the Horn of Africa. Sudan’s main indigenous cattle is the humped *Bos indicus* (zebu) type. The breed names come from human tribes, external morphological traits, specific conformation, size, and branding. The main Sudanese cattle groups include the Northern Sudan short-horned zebu (Kenana, Butana, Baggara), the Nilotic zebu of Southern Sudan (e.g. Toposa-Murle, Mangala) and the Sanga (e.g. Dinka, Nuer). Other cattle populations include White Nile cattle, Fuga or Dar EL Reeh cattle of Northern Kordofan, Nuba Mountains cattle, Gash and Aryashai of East Sudan and the Fulani cattle found in South-West Sudan and across the West Sahelian belt (Rahman 2007, Nahar 2009, HCENR 2014).

Sudan geography offers diversified agro-ecological conditions and climate, differing in rainfall, from as little as 75 mm of annual rainfall in the northern desert to about 1500 mm in the South-West forest. There is also a wide variation in the soil types, temperatures and vegetations. The north part of the country is predominantly desertic, comprising part of the Libyan desert to the West and the Nubian desert to the East, separated by a stretch of the Nile Valley. With virtually no rainfall in this region, the primary water sources that support human settlements and vegetation are a few oases in the Libyan desert. In contrast, the South and West regions consist mainly of sandy plains interrupted by mountains. It extends from the Nuba Mountains to the borders with the Central African Republic and Chad. The Northeastern parts of the country experience little rainfall and are classified as semi-desertic (Impiglia 2017). Therefore, the grazing lands in the arid and semi-arid Sudan regions are characterised by extreme temperatures, high aridity and feed and water scarcity (Musa *et al*. 2006, Ahmed *et al*., 2016). Undoubtedly, the Sudanese cattle are among the few indigenous African cattle living in extreme dryland climatic conditions. It is expected that such environmental pressures would have left selection footprints in their genomes.

The present study involves whole-genome re-sequencing of six indigenous African zebu breeds sampled in Sudan (Kenana, Butana, Aryashai, Baggara, Gash, and Fulani). Kenana is found in the region between the White and Blue Nile. Butana is located mainly in the Butana plain within a triangle delimited by the River Atbara, the Blue Nile and the River Nile. The natural habitat of Baggara cattle in Sudan is the savannah belt lying between the White Nile and the western fringes of Sudan. It is a breed associated with the Baggara Arabs ethnic group inhabiting the Sahel (mainly between Lake Chad and southern Kordofan). The Baggara nomadic pastoralists in Darfur and Kordofan moved their animals across hundreds of kilometres in search of pastures and water. Aryashai and Gash cattle are mainly found in the El Gash delta, stretching from Aroma in the South, Dordeib in the North and spreading to the Atbara River. Fulani cattle in Sudan are found in the western part of the country, in the Darfur and Blue Nile states (Figure 1). The Falata Umbororo tribes mostly own Fulani cattle in Sudan (Nahar 2009). The breed is also associated with the Fulani people living along the Sahelian belt. In addition, the Kenana and Butana breeds are anecdotally referred to as African dairy zebu because of their superior milk production capability compared to most African zebu breeds. Hence, they are mostly kept under the high input systems of dairy production (Rahman 2007). The production of F1’s heifer with exotic taurines for milk production is common across Sudanese cattle, especially among the dairy Butana and Kenana. Therefore, there is a genuine concern about the ongoing erosion of the genetic diversity of these indigenous breeds under such indiscriminate crossbreeding systems (FAO, 2009; HCENR. 2014, FAO. 2015).

**Figure 1.**
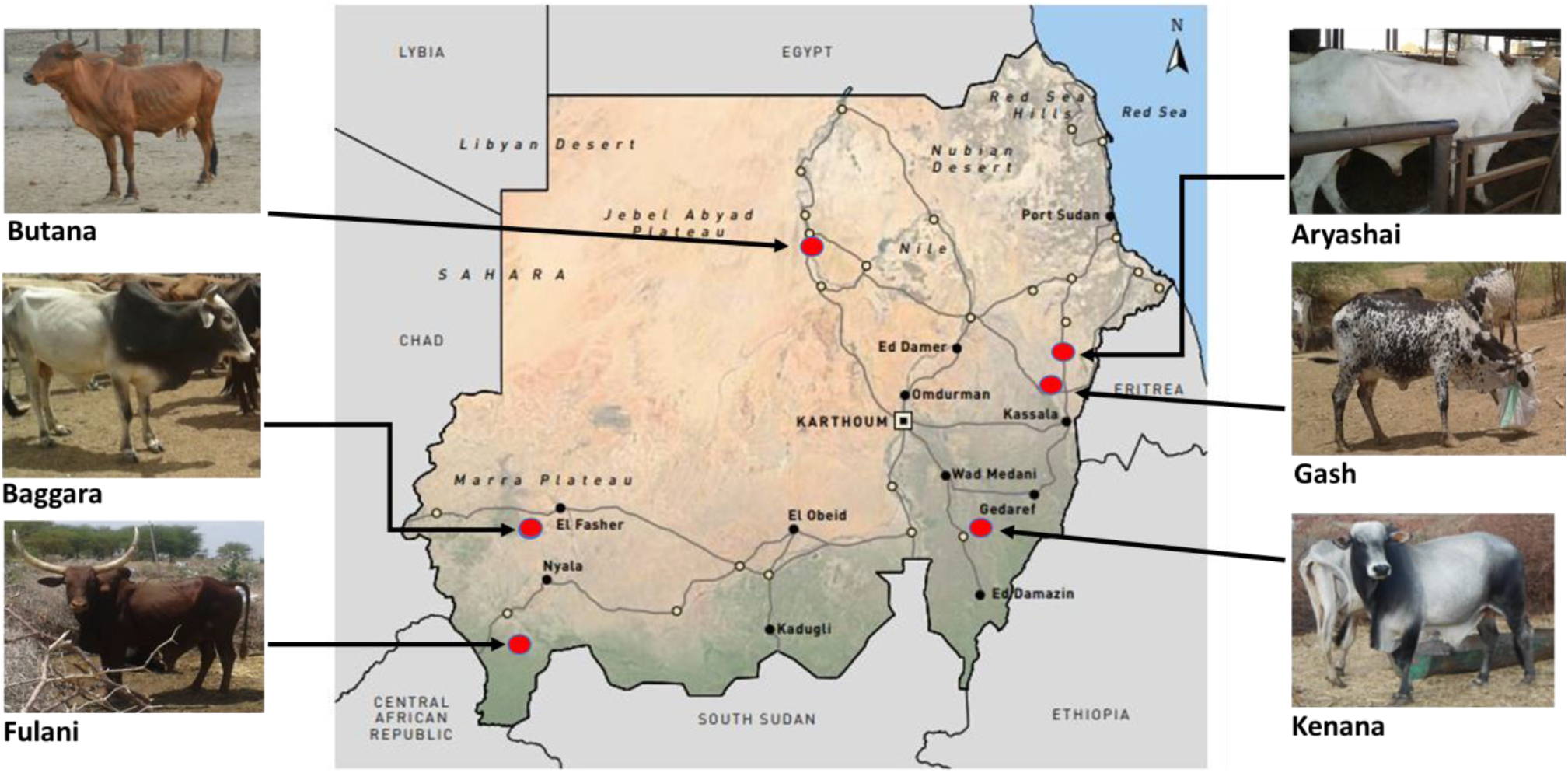
The geography of Sudan shows the sampling areas of the Sudanese zebu cattle populations. Map adapted from Impiglia (2017). Photo credit: Professor Hassan Musa, University of Khartoum, Sudan.

This study assessed the genetic diversity, population structure, and genomic footprints of positive selection in the indigenous Sudanese cattle population. In particular, we aimed to investigate the adaptation of these populations to the extreme dryland environmental condition of the African continent, addressing the question of the genetic origin, within or from outside the African continent, of their ecological adaptations. Accordingly, we applied three genomic scan approaches to detect regions of low within breed diversity (*ZH_p_*, Rubin *et al*., 2010), population divergence (*F*_ST_, Beaumont 2005) and increased haplotype homozygosity (XP-EHH, Sabeti *et al*., 2007). We performed these assessments in comparison to other cattle breeds, including two East African zebu breeds (Kenyan Boran, Ethiopian Ogaden), one African sanga breed (Ankole), one Asian zebu from Brazil (Gir), two African taurine breeds (N’Dama, Muturu), three Eurasian breeds (Eastern Finncattle, Western Finncattle and Yakutian cattle) as well two West Europe taurine breeds (Holstein and Angus).

## Materials and Methods

### Study population and sample re-sequencing

A total of 17 zebu and taurine cattle breeds comprising 185 individuals were included in the study (Table S1). We newly generated the full genome sequences of ninety indigenous Sudanese zebu cattle belonging to two independent datasets of six (Group A, *n* =60) and three (Group B, *n* = 30) zebu breeds sampled in Sudan. Group A datasets consist of 60 sequences of Kenana (KEN, *n* = 10), Butana (BTN, *n* = 10), Aryashai (ARY, *n* = 10), Baggara (BGR, *n* = 10), Gash (GAS, *n* = 10) and Fulani (FLN, *n* = 10). The Group B datasets consist of 30 sequences of Kenana (*n* = 9), Butana (*n* = 11) and Baggara (*n* =10). The 30 sequences of the Group B Sudanese datasets were generated more recently as part of the Genomic Reference Resource for African Cattle (GRRFAC) Initiative (https://grrfac.ilri.org). Therefore, they were not part of the initial variant calling described below. However, we provide additional Materials and Methods information in Supplementary file one regarding the sampling procedure, read mapping and variant calling involving the Group B dataset.

We also included new sequences of 10 Gir cattle from Brazil (GIR, *n* = 10), provided by Dr Marcos Vinicius Barbosa da Silva of Embrapa Dairy Cattle. The remaining 85 genome sequences are from the public database; Ogaden (OGD *n* = 9, Ethiopia), Kenyan Boran (BOR *n* = 10, Kenya), Savannah Muturu (MUT *n* = 10, Nigeria), N’Dama (NDM *n* = 10, The Gambia), Ankole (ANK, *n* =10, Uganda), Angus (ANG, n = 10), Holstein (HOL, *n* = 11), Eastern Finncattle (EFN *n* = 5) Western Finncattle (WFN *n* = 5) and Yakutian (YKT *n* = 5) (Stothard *et al*. 2011, Bahbahani *et al*. 2017, Kim *et al*. 2017, Weldenegodguad *et al*., 2019) The NCBI SRA accession numbers are provided in Table S1.

The ethical committee of the Faculty of Veterinary Sciences, University of Nyala, Sudan, approved the sampling protocol for the Group A Sudanese cattle. We obtained 10 ml of whole blood from each animal into EDTA VACUETTE^®^ tubes following the standard procedure under veterinarian supervision. According to the manufacturer protocol, genomic DNA was extracted from the blood samples using the Qiagen DNeasy extraction kit (Qiagen, Valencia, CA, USA). Genomic DNA was evaluated using a NanoDrop spectrophotometer (NanoDrop Technologies, USA) and gel electrophoresis. Paired-end sequencing was performed on the Illumina^®^ HiSeq platform, with the read length of PE150 bp at each end.

The Gir cattle samples were part of the progeny test program from the National Program for Improvement of Dairy Gir (PNMGL), headed by Embrapa Dairy Cattle (Juiz de Fora, Minas Gerais, Brazil) in cooperation with the Brazilian Association of Dairy Gir Breeders (ABCGIL) and the Brazilian Association of Zebu Breeders (ABCZ). Semen samples were collected for DNA extraction. For this purpose, the semen samples were washed with a lysis buffer and incubated for two hours with an extraction buffer containing dithiothreitol 10% and RNase. Pellets were incubated overnight with a saline-proteinase K buffer, and a phenol-chloroform extraction removed the proteins. The quality and quantity of DNA for all samples were evaluated using a NanoDrop 1000 spectrophotometer (Thermo Scientific, Wilmington, DE, USA). Mate-paired and paired-end libraries (200 bp and 2 x 100 bp, respectively) with different insert sizes were prepared according to the Illumina protocol and subsequently sequenced on the Illumina^®^ HiSeq platform (Illumina Inc., San Diego, CA, USA).

### Reads alignment, variants discovery and quality control

We subjected individual sample raw reads to initial quality control using Trimmomatic *v*0.38 (Bolger *et al*., 2014). First, we trimmed paired reads of adapter, low-quality bases (qscore < 20) at the beginning and end, then filtered out reads with mean q score less than 20 or length less than 35 bp. The final quality of the resulting clean reads was confirmed using FastQC v0.11.5 (https://www.bioinformatics.babraham.ac.uk/projects/fastqc/). Next, trimmed sequences were aligned to the *Bos taurus* reference genome ARS-UCD1.2 (Rosen *et al*., 2020), using BWA-MEM v0.7.17 (Li & Durbin, 2010) with default parameters. After mapping, we processed the resultant alignment files and performed variants (SNPs and InDels) discovery following the Genome Analysis Toolkit best practices pipeline (https://gatk.broadinstitute.org/hc/en-us/sections/360007226651-Best-Practices-Workflows).

Here, SAMtools *ver* 1.9 was used to convert SAM files to BAM format and for sorting by contigs (Li *et al*., 2009), duplicates were marked using Picard tools *ver* 2.18.2 (http://broadinstitute.github.io/picard/). Variant calling was performed on individual cattle samples using the Haplotype caller of GATK v3.8-1-0-gf15c1c3ef (McKenna *et al*., 2010) and incorporating known variants from dbSNP *ver*150 (Cunningham *et al*., 2019). The GATK joint genotyping approach (*GenotypeGVCFs* mode) was adopted to simultaneously identify variants in all cattle samples.

We subjected SNPs and insertions/deletions (InDels) separately to the GATK hard filter (VariantFiltration) steps. The filtering criteria for SNPs include (QD > 2.0, MQ > 40, ReadPosRankSum > 8.0, HaplotypeScore > 13, MappingQualityRankSum > 12.5). The autosomal biallelic SNPs which passed the above filtering criteria and having a Phred-scaled quality score of above 20 (QUAL >20; approximately 99% likelihood of being correct) were retained for further analyses. The filter criteria for detected InDels include (QD < 2.0 || FS > 200.0 || ReadPosRankSum < −20.0 || QUAL < 20).

Furthermore, the proportion of missing genotypes in an individual sample and the relatedness among other samples were estimated using the options “--*missing-indv*” and “—*relatedness*”, respectively, of Vcftools *v0.1.15* (Danecek *et al*., 2011). Both individuals of a pair of samples with high relatedness (> 0.8) were excluded from the dataset if they belong to different breeds. In contrast, only one of two individuals with high relatedness was excluded if they belonged to the same breed. Following these criteria, five samples, including two Butana (*n* = 2), two Gash (*n* =2) and one Baggara (*n* = 1), were removed. No animal was removed due to excessive missing data (> 20%). Hence, 150 cattle samples were retained in downstream analyses.

The total numbers of variants (SNPs and InDels) and the transition to transversion (Ts/Tv) ratio for individual cattle breeds were estimated using BCftools *ver* 1.8 (Li *et al*., 2009). SnpEff v4.3t (Cingolani *et al*., 2012) was used to ascertain the genome location and effects of detected variants based on the *Ensembl* cow gene database (ARS-UCD1.2) dbSNP *ver*150. The proportion of detected variants was classified as “known” if the non-reference allele is present in the dbSNP, otherwise as “novel.”

### Genomic diversity and inbreeding

From the detected high-quality autosomal SNPs (~ 42.9 M), we estimated the counts of segregating sites for individual cattle using PLINK v2 (Purcell *et al*., 2007). The average of individuals of the same breed was reported as the per breed estimate. Within-population nucleotide diversity (π) values (Nei and Li 1979) and global averages of pairwise population differentiation (*F*_ST_) (Weir and Cockerham 1984) among cattle breeds were estimated based on overlapping 100 kb window and 50 kb step size along bovine autosomes using Vcftools *v0.1.15*. Due to the differences in the number of samples in some of the studied breeds, we randomly selected five representative samples from each breed to estimate these latter parameters.

### Population genetic structure

We performed Principal Component Analysis (PCA), admixture, and maximum likelihood tree (Treemix) to investigate the genetic structure among the different cattle breeds using whole-genome autosomal SNPs. Linkage disequilibrium (LD) pruning of SNPs was performed using PLINK default option “50 Kb step 10Kb SNPs, r^2^ > 0.5” (Purcell *et al*., 2007), resulting in approximately 24 million SNPs. We further used PLINK to generate PCA eigenvectors and eigenvalues in two categories: all 17 cattle breeds and Sudanese zebu samples only. PCA plots were generated for the first two eigenvectors using the ggplot2 package in the R *ver* 3.6.3 environment.

The block relaxation algorithm implemented in the ADMIXTURE *ver 1.3.0* software (Alexander et al., 2009) was used to estimate the ancestry proportions of individual cattle samples with kinship (K) set from 2 to 5. Admixture analysis was preceded by removing SNPs with more than 10% missing data (--*geno 0.1*) using VCFtools. Next, we selected the optimal K based on the cross-validation error procedure. Finally, the admixture ancestry ratio was visualised using the pophelper R package (Francis, 2017).

Phylogenetic relationships based on maximum likelihood tree and possible gene flow events among cattle breeds were investigated using Treemix *v*1.3 (Pickrell & Pritchard, 2012). The phylogenetic tree was constructed several times, incorporating possible migratory events, from m0 (no migration event) to m10 (ten migration events). Finally, the models were visualised in R using the script provided in Treemix.

### Genome diversity analysis and functional annotation

We applied three genomic scan approaches to detect regions of low within breed diversity (*ZHp*), population differentiation (*F*_ST_) and increased haplotype homozygosity (XP-EHH) as a proxy of signals for candidate selective sweep in the indigenous African zebu from Sudan.

We applied the within-population *Hp* test to detect genomic signatures of low diversity in each of the eight African zebu breeds and the Gir breed. The *Hp* analysis involves counting reads with the most and most minor abundantly observed alleles at every SNP position within a specified window size and sliding step size. The distribution of *Hp* values was normalised by transforming *Hp* to Z-scores (*ZHp*) using (*ZHp* = (*Hp* - *μHp*)/*σHp*) (Rubin *et al*., 2010, Rubin *et al*., 2012, Axelsson *et al*., 2013). Using VCFtools (Danecek *et al*., 2011), we estimated regions of population divergence (*F_ST_*) based on Weir and Cockerham (1984) between the Sudanese zebu populations and the combined 11 zebu and taurine breeds (Gir, Ogaden, Kenya Boran, Ankole, Muturu, N’Dama, Angus, Holstein, Western Finncattle, Eastern Finncattle and Yakutian). We performed two genome-wide *F*_ST_ analyses using independent datasets of Sudanese zebu populations; Groups A (Arayshai, Baggara, Butana, Gash, Fulani and Kenana) and B (Baggara, Butana and Kenana). The weighted *F_ST_* were also *Z*-transformed in R. In each *ZH*p and *F*_ST_ test, we estimated genome-wide test statistics using 100 kb windows and sliding 50 kb step across the bovine 29 autosomes and excluded windows containing less than 10 SNPs. Also, the 100-kb windows in the extreme top 0.5% test values were arbitrarily defined as candidate selection outlier windows. We find an overlap of detected selective sweep regions between the two *F*_ST_ analyses using BEDTools intersect (*version* 0.2.29) (Quinlan and Hall, 2010).

*Bos taurus* (ARS-UCD1.2) genes overlapping the candidate selected windows were retrieved based on the *Ensembl* cow genes database 104 using the *Ensembl* BioMart online tool (http://www.ensembl.org/biomart) (Smedley *et al*., 2009). Then, we generated a Venn diagram based on the list of detected selective sweep regions based on *ZHp* and *F_ST_* using the online tool at (https://molbiotools.com/listcompare.html).

Finally, following the *ZH*p and *F*_ST_ analysis results, we further investigate the candidate selective sweep on chromosome 16 uniquely detected in five of the six Sudanese zebu breeds by performing the LD-based XP-EHH test using the Rehh *ver* 2.0 R package (Gautier *et al*., 2017). Here, we contrasted the combined Group A Sudanese zebu breeds (except Fulani) against the Fulani, the other zebu breeds (Ethiopia Ogaden, Kenya Boran and Gir), and the taurine breeds (African and European taurine). As this analysis required phased haplotype, we performed phasing of the entire chromosome 16 SNP data for all 155 cattle samples (excluding Group B Sudanese sequences) using the default parameter of Beagle ver 5.0 (Browning *et al*., 2007), except for *Ne*, which was set to 1000 to improve the accuracy of phase information (Dutta *et al*., 2020)

## Results

### Sequence reads and variants statistics

After removing adapter sequences and low-quality reads, we mapped individual sequences to the *Bos taurus* genome of reference, ARS-UCD1.2 (Rosen *et al*. 2020). We achieved an average alignment rate of up to 99% in all samples. Approximately 43 million SNPs were detected in all the 150 cattle samples. However, the number of segregating sites (homozygous and heterozygous alternate SNPs) in each breed ranges from more than 23 million in the zebu (humped) breeds to between 9 and 14 million in the taurine (humpless) breeds. Within individual humped cattle, the average number of SNPs ranged from around nine million in Kenya Boran to 12 million in Kenana and Butana. On the other hand, a range of 4.5 million to 6.3 million SNPs was detected in the humpless African and European cattle. Therefore, the number of SNPs identified in individual humped cattle is approximately 2 – 2.5 times higher than the number of SNPs in individual taurine cattle (Table S2).

The estimated heterozygous to homozygous SNPs in most 17 cattle breeds is more than 1. The exceptions being the Gir (Het/Hom = 0.74), the Muturu (Het/Hom = 0.63) and the Eastern Finncattle (Het/Hom = 0.86). The highest ratio among the Sudanese zebu breeds is in Butana (Het/Hom = 1.53), while the lowest is observed in Gash (Het/Hom = 1.01). The average transition versus transversion (Ts/Tv) ratio for the 17 cattle breeds is above 2.3 (Table S2).

The numbers of small insertions and deletions (InDels) in individual humped cattle range from about 1.2 million in most African and Asian breeds to approximately 1.5 million in Kenana and Butana. In comparison, the number of InDels in the taurine individuals varies between 600,000 to 800,000 (Table S2). In contrast to the database of known cattle variants (dbSNP *ver*150, last accessed December 2020), an average of 1.7 M (~ 6.5%) novel SNPs and 1.7 M (~5 0.5%) novel InDels were detected for the Sudanese population (Table S2). These represent a valuable addition to the database of known cattle variants.

### Genetic diversity and population differentiation

The average nucleotides diversities among 17 cattle breeds studied are the highest in the eight African zebu breeds (between 2.63 x 10^-3^ and 3.2 x 10^-3^), followed by the Gir (2.49 x 10 ^-3^), the African sanga, Ankole (2.20 x 10^-3^), and the taurine breeds (between 1.50 x 10 ^-3^ to 1.24 x 10 ^-3^). The lowest value is observed in Muturu (9.94 x 10^-4^) (Figure 2). On the other hand, nucleotide diversities are the highest within the African zebu breeds in the two African zebu dairy breeds, Butana and Kenana (~ 3.2 x 10-3) (Figure 2A).

**Figure 2.**
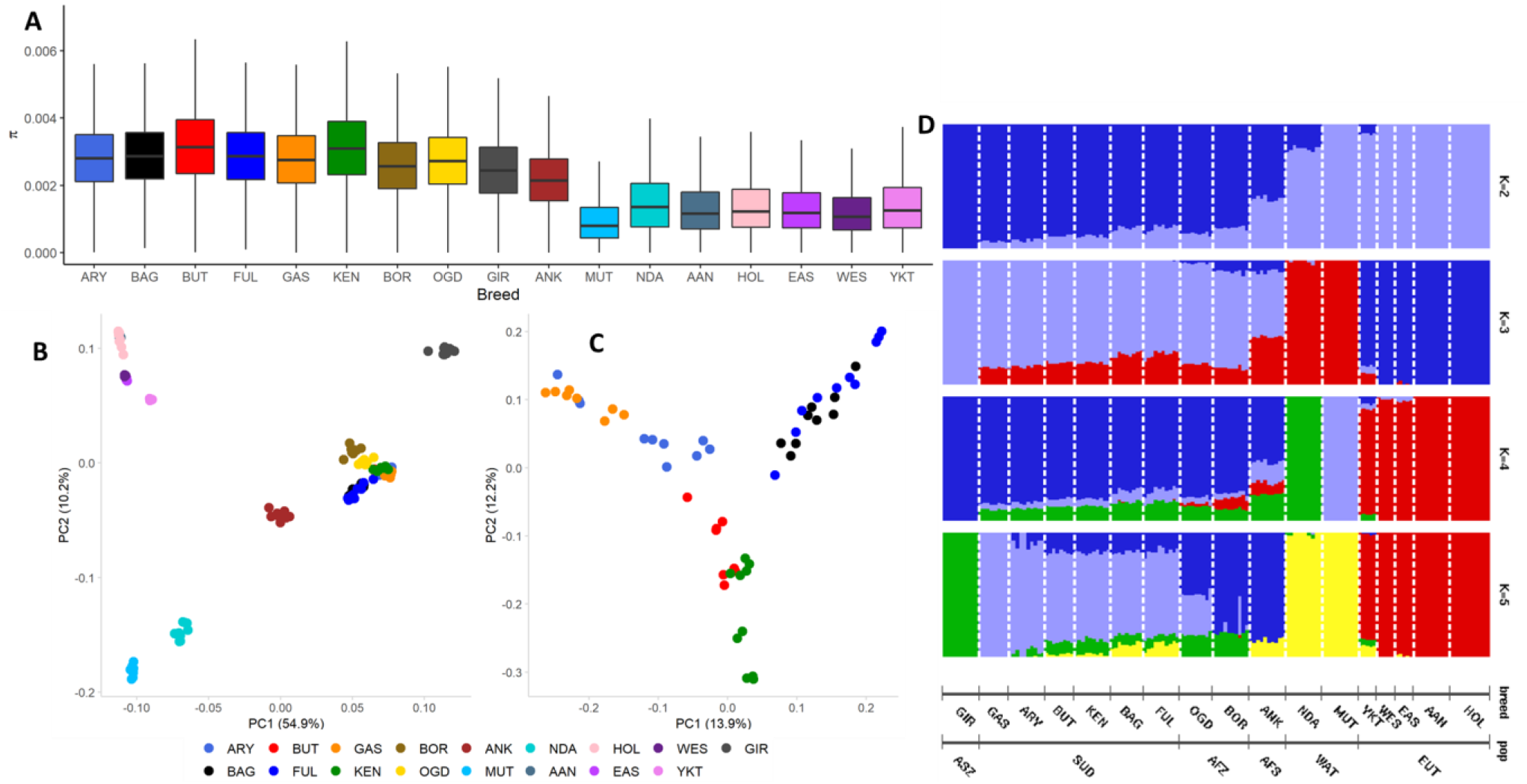
Genetic diversity and population structure of the 17 studied breeds. **(A)** Boxplot of average nucleotide diversities (π). The nucleotide diversity level within each cattle breed was calculated based on an overlapping 100 kb window with a 50 kb step size. **(B)** Principal component (PC 1 *versus* PC 2) analysis for 17 cattle breeds. **(C)** Principal component (PC 1 *versus* PC 2) analysis for the six Sudanese zebu cattle populations. **(D)** Admixture plot showing ancestry proportions for the 17 cattle breeds. The population structure was assessed using ADMIXTURE *ver*.1.3.0. The individual population are represented by a vertical bar and partitioned into coloured segments. Each segment’s length represents the proportion of the inferred number of ancestries (K = 2 to K = 5). GIR – Gir, GAS – Gash, ARY – Aryashai, BTN – Butana, KEN – Kenana, BGR – Baggara, FLN – Fulani, OGD – Ogaden, BOR – Kenya Boran, ANK – Ankole, NDA – N’Dama, MUT – Muturu, YKT – Yakutian, WES – Western Finncattle, EAS – Eastern Finncattle, AAN – Angus and HOL – Holstein. The cattle breeds were also grouped into six entities, namely, Asian zebu (ASZ), Sudanese zebu (SUD), East African zebu (AFZ), African sanga (AFS), West African taurine (WAT) and European taurine (EUT).

Population differentiation (*F*_ST_) is low among the African zebu breeds (Table S3). The average highest population divergence is between the Sudanese Gash and the taurine breeds (*F_ST_* = 0.324). It is the lowest between the Sudanese Baggara and the taurine breeds (*F_ST_* = 0.267). Interestingly, the average population divergence between the African zebu and the West African taurine is higher for the West African shorthorn, Muturu (*F_ST_* =0.342), compared to the N’Dama, a West African longhorn (*F_ST_* =0.245) (Table S3). Population divergence between each of the African zebu and the Gir cattle is around 0.1.

### Population structure

The principal component analysis of the 17 cattle breeds was performed using approximately 24 million autosomal bi-allelic SNPs after removing SNPs with a minor allele frequency of less than 0.05. The first and second principal components account for 54.9% and 12.2% variations. PC 1 clearly distinguished the taurine from the zebu breeds, while PC 2 separates African breeds from non-African breeds (Figure 2B). A second PCA, including only the Sudanese zebu, reveals pairs of clusters (Kenana - Butana, Aryashai - Gash, and Baggara – Fulani), with no clear differentiation between the Baggara and the Fulani. PC 2 (12.1%) shows the separation of Kenana and Butana (African zebu dairy breeds) from the other Sudanese zebu (Figure 2C).

After LD pruning (r^2^ > 0.5), a dataset of approximately 4.5 million SNPs was used to explore the ancestry proportions (K) of the different cattle breeds for K ranging from 2 - 5 (Figure 2D). The admixture cross-validation (CV) option (Alexander *et al*. 2009) supports K = 4 with the lowest CV error as the likely optimum number of ancestral backgrounds for our populations. The four possible ancestral backgrounds inferred here are *Bos indicus*, European *Bos taurus*, and two distinct African *Bos taurus* ancestries (N’Dama and Muturu). At this level, the African zebu breeds show admixed zebu and taurine genetic backgrounds, as in previous reports (Kim *et al*., 2020), including two distinct African *Bos taurus* ancestries, the longhorn N’Dama and the shorthorn Muturu (Tijjani *et al*., 2019). On the other hand, a low level of European taurine ancestral background is observed in the Sudanese Baggara, Fulani and the two East African zebu breeds (Figure 2D). Furthermore, at five ancestral backgrounds (K = 5) reveal two separate African zebu ancestries, whereby the dryland Sudanese zebu are distinct from the East African zebu breeds.

Both PC analyses (Figure 2C) and admixture analysis (Figure 2D, K = 2 to 5) indicate substantial genetic similarity between the Sudanese Baggara and the trans-Sahelian Fulani, with no separation between the two populations in the PC analysis and very similar ancestral background (admixture analysis).

#### Evolutionary relationships among cattle breeds

As for the admixture analyses, genome-wide unlinked autosomal SNPs were used for the Treemix analyses. The phylogenetic relationships and migration events reveal several possible genes flows among the cattle breeds (Figure 3). By sequentially adding up to 10 migration events and agreeing with the admixture analysis, we observe possible gene flow from the African taurine into most African zebu breeds, including four Sudanese populations (Kenana, Butana, Baggara and Fulani). On the other hand, we did not observe gene flow between the Sudanese zebu and any European breeds. Our Treemix results also reveal possible introgression between the two West African taurine breeds with the direction of gene flow from the shorthorn Muturu to the longhorn N’Dama (Figure 3).

**Figure 3.**
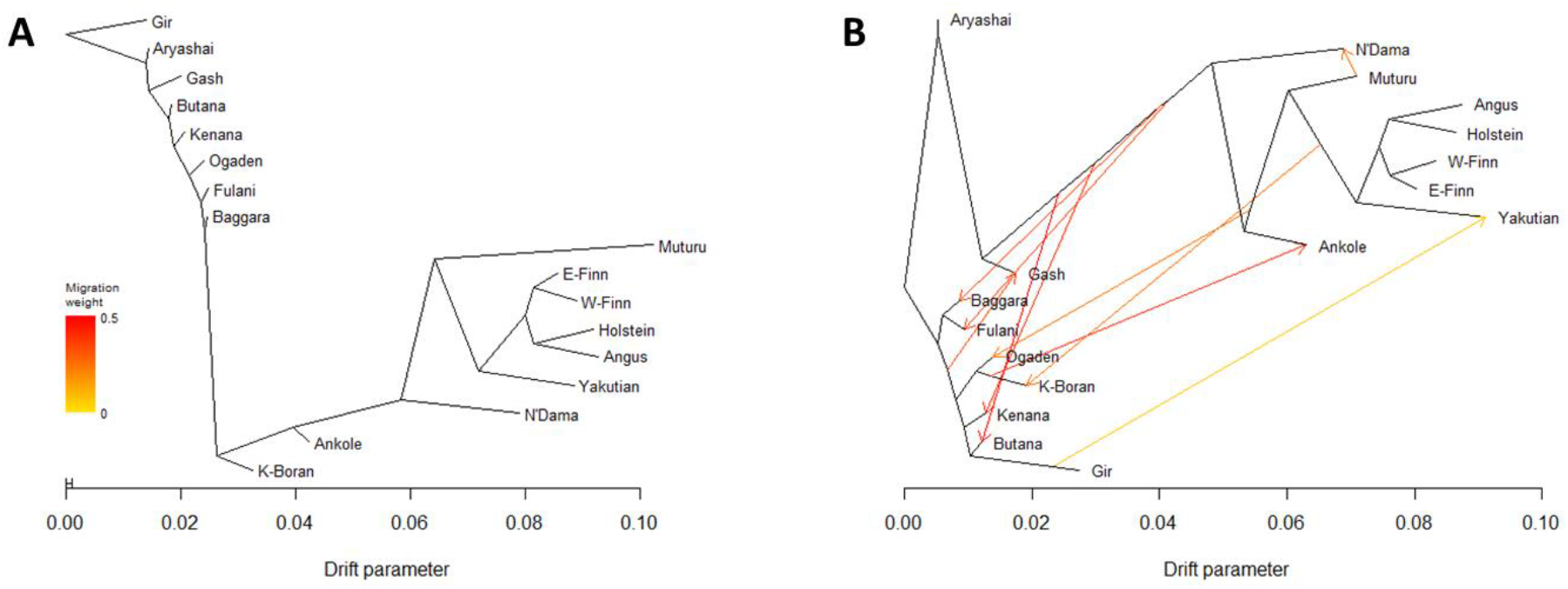
Maximum likelihood evolutionary tree and possible gene flow among the 17 cattle breeds: **(A)** without migration events, **(B)** assuming ten migration events. E-Finn Eastern Finncattle) W-Finn r(Western Finncattle), K-Boran (Kenyan Boran).

### Genomic signatures of positive selection in zebu populations

We performed a genome-wide autosomal *Hp* selection scan in the nine zebu breeds of six Sudanese zebu populations, Kenyan Boran, Ogaden and Gir. The distributions of *Z*-transformed *Hp* scores across the bovine autosome are shown in Figure 4 (Sudanese populations) and Figure S1 (other zebu breeds). We considered 249 outlier windows (100-kb windows in the lowest 0.5% of *ZHp* scores) in each breed as regions with significantly reduced diversity and possible signatures of positive selection (Tables S4 - S12). The highest *ZHp* threshold is observed in Baggara (*ZHp* = −3.251), while the lowest is observed in Gir (*ZHp* = −3.958) (Figures 4 and S1). We identify 273 outlier windows common to at least two breeds among the Sudanese zebu population. We merged the windows in the proximity of up to 5 kb, resulting in 114 selective sweeps regions varying in size from 0.1 to 1.5 Mb. Fifty-five selective sweep regions overlap with outlier windows detected in at least one non-Sudanese zebu population. The remaining 59 selective sweep regions were detected exclusively in the Sudanese zebu populations. Thus, these regions are shared and unique Sudanese zebu selective sweep regions (Table S13). Based on *Ensembl* cow genes 104 (ARS-UCD1.2), 47 out of the 55 shared Sudanese – non-Sudanese zebu selective sweep regions overlap with protein-coding genes (*n* = 227) (Tables S14). In contrast, of the 59 unique Sudanese selective sweeps, 40 overlaps with 135 protein-coding genes (Tables S15). In both cases, more than 60 per cent of the annotated regions contain more than one protein-coding gene (gene-rich). However, the selective sweep regions devoid of any annotated cow genes (desert gene regions) spans a total 4.15 Mb ARS-UCD1.2 genome region (Table S16)

**Figure 4.**
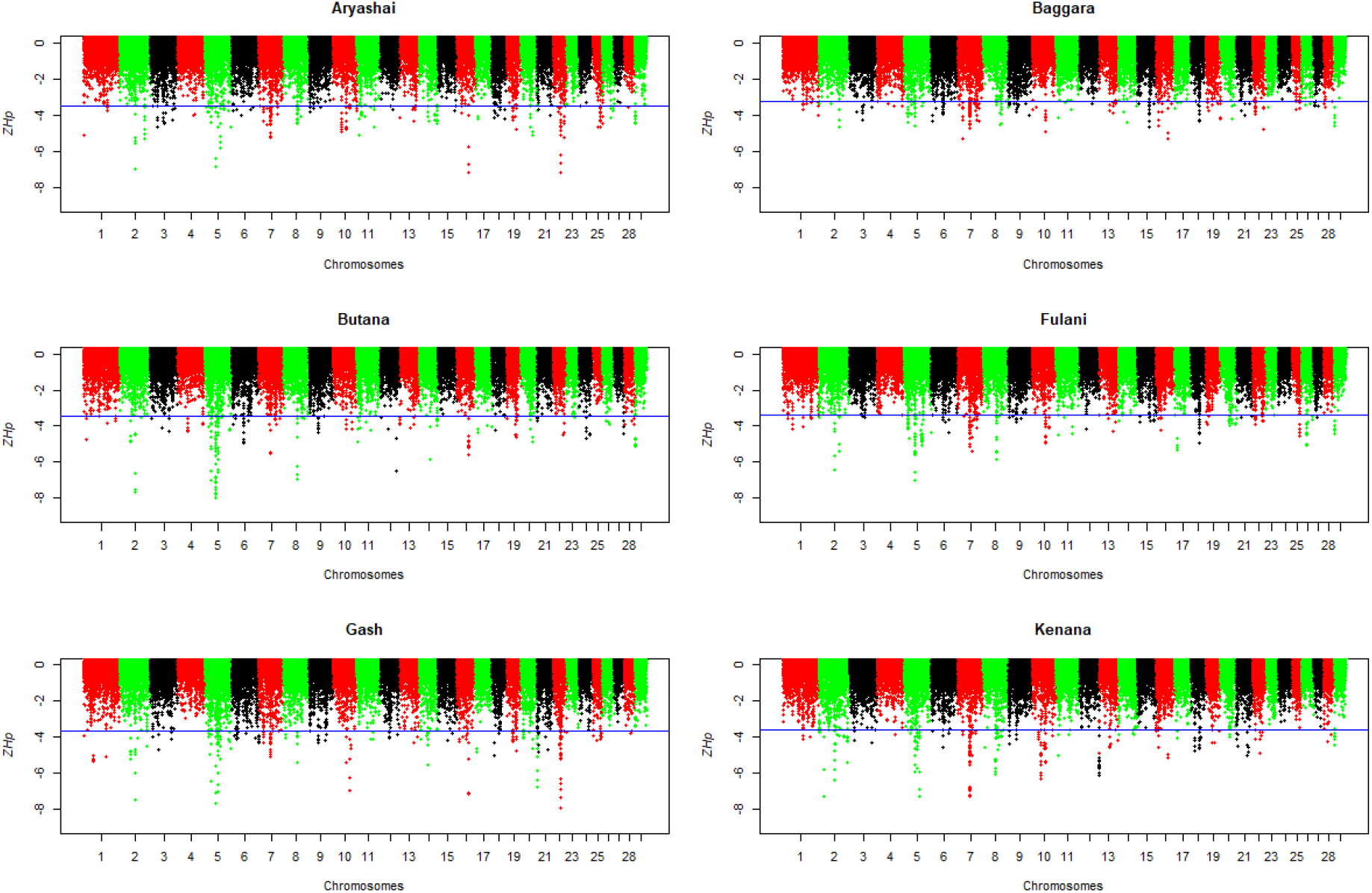
Genome-wide distribution of *ZHp* scores across bovine autosomes in the six Sudanese zebu populations. The blue line indicates the *ZHp* threshold value (lowest 0.5%) for selecting outlier windows (candidate regions under positive selection).

### Shared African and Asian zebu selective sweep regions

Among the 47 shared selective sweep regions overlapping annotated genes, two regions, on chromosome 5 (5:47.40 – 48.0 Mb) and 7 (7:49.85 – 51.15 Mb), are detected in eight of the nine zebu breeds. We detect the 600 kb region on chromosome 5 in all the six Sudanese zebu populations, the Kenyan Boran and Gir breeds but not in the Ethiopian Ogaden. The region overlaps eight genes (*ENSBTAG00000052954*, *ENSBTAG00000053419*, *GRIP1*, *HELB*, *HMGA2*, *IRAK3*, *LLPH*, *TMBIM4*) with genes function related to growth, conformation, reproduction and the immune systems (Porto-Neto *et al*., 2014, Naval-Sánchez *et al*., 2020). The approximately ~1.5 Mb gene-rich sweep on chromosome 7 overlaps with up to 28 protein-coding genes, including two heat shock proteins, *DNAJC18* and *HSPA9*. Again, this large sweep is detected in eight zebu breeds, except the Sudanese Butana. Other shared selective sweep regions between African and Asian breeds (present in at least six breeds) are found on chromosomes 2 (2:70.25 - 70.50 Mb), 8 (8:59.35 - 59.70 Mb), 10 (10:58.85 - 59.15 Mb) and 17 (17:13.15 - 13.35 Mb) (Table S14).

We found several candidates selected regions in African zebu breeds which are not present in the Asian Gir. These include regions on chromosomes 7 (7:52.10 – 52.40 Mb), 11 (11:13.10 – 13.30 Mb), and 20 (20:48.80 – 48.95 Mb) detected in seven of the eight African breeds. The exceptions are Butana (chromosome 7), Gash (chromosome 11) and Kenana (chromosome 20) (Table S14). In particular, the region on chromosome 7 overlaps with up to eight protein-coding genes (*ENSBTAG00000030474*, *ENSBTAG00000049316*, *ENSBTAG00000049493*, *ENSBTAG00000050840*, *ENSBTAG00000053160*, *PCDHAC2*, *PCDHB1* and *SLC25A2*). Also, we found another gene-rich region on chromosome 19 (19:26.35 – 26.50 Mb), containing up to fourteen protein-coding genes detected in six African zebu breeds, including four Sudanese populations (Table S14).

Besides, we identified regions in the Sudanese zebu population shared only with the Gir. These include a 650 kb region on chromosome 5 (5:48.4 - 49.05 Mb) found in five Sudanese zebu populations, the exception being the Baggara. This region overlaps five genes (*ENSBTAG00000000237*, *LEMD3*, *MSRB3*, *TBC1D30* and *WIF1*). Also, a 150 kb candidate selected region on chromosome 2 is detected in four Sudanese populations and the Gir. It overlaps with the macrophage receptor with the collagenous structure gene (*MARCO*), involved in the innate immune response (Kissick *et al*., 2014). Additionally, a 150 kb region on chromosome 5, overlapping with the *HOXC* gene cluster, was also detected exclusively in four Sudanese zebu populations and the Gir (Table S14). This region has also been reported previously in the Gir cattle (Liao *et al*., 2013).

### Sudanese zebu specific candidate selective sweeps

We classified as Sudanese zebu-specific sweeps 59 genome regions of low diversity (*ZHp*) detected uniquely in the Sudanese zebu populations. The commonest sweep regions are a 450 kb region on chromosome 7 (7:54.0 – 54.45 Mb) and a 250 kb region on chromosome 16 (16: 50.40 – 50.65 Mb). They are present in five Sudanese breeds, the exception being Baggara and Fulani, respectively (Table S15). The region on chromosome 7 contains *ARHGAP26* and *NR3C1*, while chromosome 16 region contains seven genes (*FAAP20*, *MORN1*, *PEX10*, *PLCH2*, *PRKCZ*, *RER1*, and *SKI*). In addition, we identified in four Sudanese breeds a 200 kb region on chromosome 7 (7:19.85 – 20.05 Mb). This region is downstream of another 150 kb region (7:19.60 – 19.75 Mb) detected in three Sudanese populations. The two regions are close to each other, being only separated by 100 kb. Together, they overlap with 16 genes (Table S15). More so, a gene-rich region of 200 kb on chromosome 5 (5: 55.85 - 56.05 Mb) was detected in three populations. It includes seventeen genes (*ARHGAP9*, *ARHGEF25*, *B4GALNT1*, *DCTN2*, *DDIT3*, *DTX3*, *ENSBTAG00000049386*, *ENSBTAG00000051574*, *ENSBTAG00000051593*, *GLI1, INHBC*, *INHBE*, *KIF5A*, *MARS1*, *MBD6*, *PIP4K2C*, and *SLC26A10*). The remaining regions overlapping at least one protein-coding gene are detected only in two Sudanese zebu populations (Table S15).

### Genetic differentiation at shared and unique Sudanese zebu candidate selected genome regions

We searched for selective sweeps possibly linked to environmental adaptation in the arid and semi-arid African regions. Hence, we looked for evidence of divergence at genome regions between the Sudanese zebu population and the other studied cattle populations following the Weir and Cockerham (1984) genetic differentiation index (*F_ST_*). Firstly, we calculated *F_ST_* values between the combined six Sudanese zebu populations and the combined 11 non-Sudanese zebu and taurine breeds (see Materials and Methods). The genome-wide distribution of Z-transformed *F_ST_* values across the 29 bovine autosomes is presented in Figure 5A. We identified 106 outliers 100-kb windows in the top 0.5 % positive *ZF_ST_* values (0.3 ≤ *F*_ST_ ≤ 0.5 and 4.1 ≤ *ZF*_ST_ ≤ 7.4). Neighbouring windows were merged, resulting in 33 selective sweeps spanning 149 protein-coding genes (Table S17).

**Figure 5.**
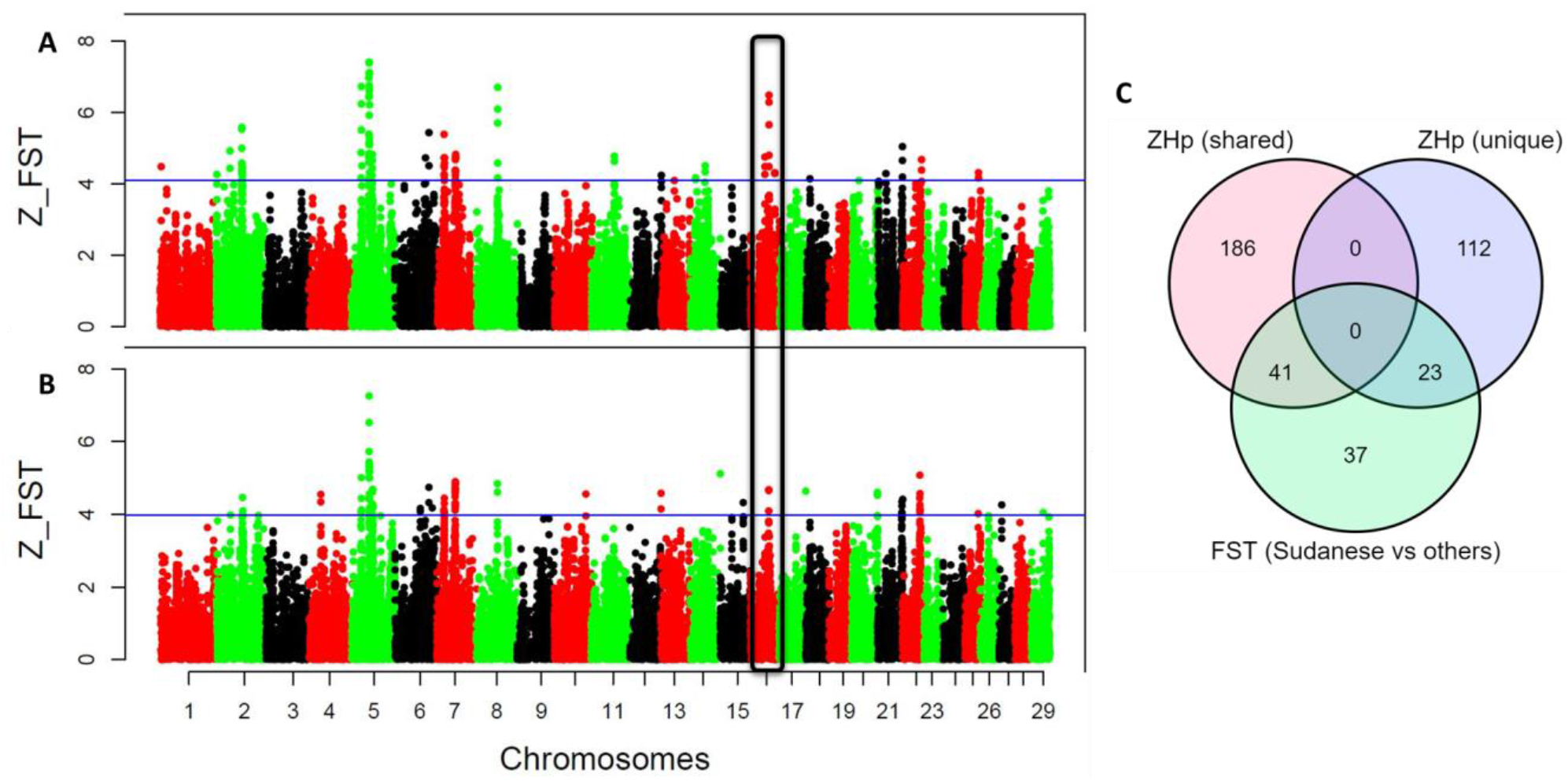
Genome-wide distribution of *ZF*_ST_ scores along *Bos taurus* (ARS-UCD1.2) autosomes following the population differentiation *F*_ST_ analyses between each of two different datasets of Sudanese zebu populations **(A).**Group A, *n* = 60 and **(B).** Group B, *n* = 30, combined eleven other zebu and taurine cattle breeds. The Group A populations consist of six Sudanese breeds, namely; Aryashia, Baggara, Butana, Fulani, Gash and Kanana. Group B samples consist of Baggara, Butana and Kenana breeds. The blue lines indicate the top 0.5% *ZF*_ST_ values threshold to consider outlier regions. **(C).** Venn diagram showing the numbers of overlapping detected protein-coding genes between *ZH*p and *F*_ST_ tests. The *F*_ST_ detected genes are the 101 common genes following the two *F*_ST_ analyses involving group A and group B Sudanese zebu population

Next, we repeated the genome-wide *F_ST_* analysis with an independent dataset of Sudanese zebu cattle genome sequences to further confirm the selected regions of population divergence between the indigenous Sudanese zebu population and other non-Sudanese cattle populations. We refer to the initial Sudanese genome dataset as Group A and the new genome dataset as Group B. The new dataset comprises 30 newly generated genome sequences of Baggara (*n* = 10), Butana (*n* = 11) and Kenana (*n* = 9) from the Genomic Reference Resource for Africa cattle (GRRFAC) initiative. Following a similar *F_ST_* approach with the Group A dataset, we identified 105 outliers 100-kb windows within the top 0.5 % positive *ZF*_ST_ values (~ ≥ 0.3 *F*_ST_ ≤ 0.5 and ≥ 3.9 *ZF*_ST_ ≤ 7.25). These windows were merged into 36 selective sweep regions overlapping 154 protein-coding genes (Figure 5B, Table S18).

By comparing the two *F*_ST_ results, we found an overlap of 101 protein-coding, positively selected genes clustered into eighteen mostly gene-rich selective sweep regions (Table S19). Among them are 11 selective sweeps regions classified as shared (*n* = 8) and unique (*n* = 3) Sudanese zebu selective sweep spanning 41 and 23 positive selected genes, respectively (Figure 5C, Table 1). Of the eight highly differentiated, shared selective sweep regions, four are found on chromosome 5 (*n* = 4). In addition, the strongest *F*_ST_ signals in the two analyses (mean *Z*_*F*_ST_ = 7.18) are found on one of the regions on chromosome 5 (5:47.25 – 47.95) (Figure 5A-B). The remaining highly differentiated and shared selective sweeps are found on chromosomes 2 (*n* = 1), 7 (n = 2) and 8 (*n* = 1) (Tables 1). The two regions on chromosome 7 add up to 0.8 Mb and are part of the *ZHp* detected 1.5 Mb region (Table S14)

**Table 1.** Examples of shared with non-Sudanese and Sudanese specific gene-rich and high population divergent selection signatures of putatively linked adaptive phenotypes for the dryland environment.

Genomic signatures of desert adaptation
| Chr | Start<sup>\$</sup> | End<sup>\$</sup> | ZHp | F<sub>ST</sub><sup>*</sup> | ZF<sub>ST</sub> | Overlapping *Bos taurus* genes<sup>^</sup> |
| --- | --- | --- | --- | --- | --- | --- |
| 2 | 70.60 | 70.80 | Shared | 0.42 | 5.02 | *EN1, MARCO* |
| 5 | 25.85 | 26.10 | Shared | 0.47 | 5.87 | *HOXC10,HOXC11,HOXC12,HOXC13,HOXC4,HOXC5,HOXC6,HOXC8,HOXC9,SMUG1* |
| 5 | 47.25 | 47.95 | Shared | 0.55 | 7.18 | *ENSBTAG00000052954,ENSBTAG00000053419,GRIP1,HELB,HMGA2,IRAK3,LLPH,TMBIM4* |
| 5 | 48.35 | 48.85 | Shared | 0.50 | 6.38 | *LEMD3,MSRB3,WIF1* |
| 5 | 55.85 | 55.95 | Unique | 0.30 | 4.2 | *ARHGEF25, B4GALNT1, DCTN2, DTX3, ENSBTAG00000049386, ENSBTAG00000051574, ENSBTAG00000051593, KIF5A, MBD6, PIP4K2C, SLC26A10* |
| 5 | 56.90 | 57.10 | Shared | 0.40 | 4.76 | *ANKRD52,APOF,APON,CNPY2,COQ10A,CS,ENSBTAG00000052361,GLS2,IL23A,MIP,NABP2,PAN2,SLC39A5,SPRYD4,STAT2,TIMELESS* |
| 7 | 19.75 | 20.00 | Unique | 0.40 | 4.5 | *CREB3L3,MAP2K2,PIAS4,SIRT6,STAP2,ZBTB7A* |
| 7 | 50.00 | 50.20 | Shared | 0.39 | 4.49 | *CTNNA1,ENSBTAG00000004415,LRRTM2* |
| 7 | 50.60 | 51.10 | Shared | 0.41 | 4.86 | *CXXC5,DNAJC18,ECSCR,MZB1,NRG2,PROB1,PSD2,SLC23A1,SMIM33,SPATA24,STING1,UBE2D2* |
| 8 | 57.80 | 58.05 | Shared | 0.46 | 5.78 | *ENSBTAG00000005667* |
| 16 | 50.40 | 50.65 | Unique | 0.45 | 5.58 | *FAAP20, MORN1, PEX10, PLCH2, PRKCZ, RERI, SKI* |

<sup>\$</sup>Unit is in megabases (Mb)
<sup>\*</sup>population differentiation ( $F_{ST}$ ) between combined Sudanese zebu breeds and the other eleven zebu and taurine cattle breeds studied.
<sup>^</sup>Positive selected genes based on the overlap between ZHp and $F_{ST}$ genomic scan tests in Sudanese zebu breeds.

The 23 Sudanese zebu-specific highly differentiated, positive selected genes are distributed across three selective sweep regions on chromosomes 5, 7 and 16. Each of these regions spans 11, 5 and 7 protein-coding genes, respectively (Table 1). Intriguingly, at least one gene within these regions is directly or indirectly involved in insulin signalling and glucose metabolism, which are likely relevant adaptive metabolic processes in hot and arid environments (Rocha *et al*., 2021). Examples of these genes are *KIF5A* (Cui *et al*., 2011), *PIP4K2C* (Shim *et al*., 2016), *MAP2K2* (Laskowski *et al*., 2016), *FAAP20* (Lagundžin *et al*., 2019), *PRKCZ* (Zou *et al*., 2013, Ahmed *et al*., 2020) and *SKI* (Diaz *et al*., 2012). However, while the chromosomes 5 and 7 unique selective sweeps are detected in fewer Sudanese zebu breeds (≤ 4) than the chromosome 16 region detected exclusively in five Sudanese breeds (except the Fulani). Also, the chromosome 16 region shows the strongest average *F*_ST_ signal (Tables S15 and 1). Likewise, this region is highly differentiated between the five Sudanese zebu and the taurine populations (combined European and African taurine breeds) (*F*_ST_ > ~0.75), non-Sudanese zebu populations (East African and Asian zebu) and Sudanese Fulani (~ ≥ 0.1 *F*_ST_ ≤ 0.25) (Figure 6B). Hence, we opted to investigate this particular region further.

**Figure 6.**
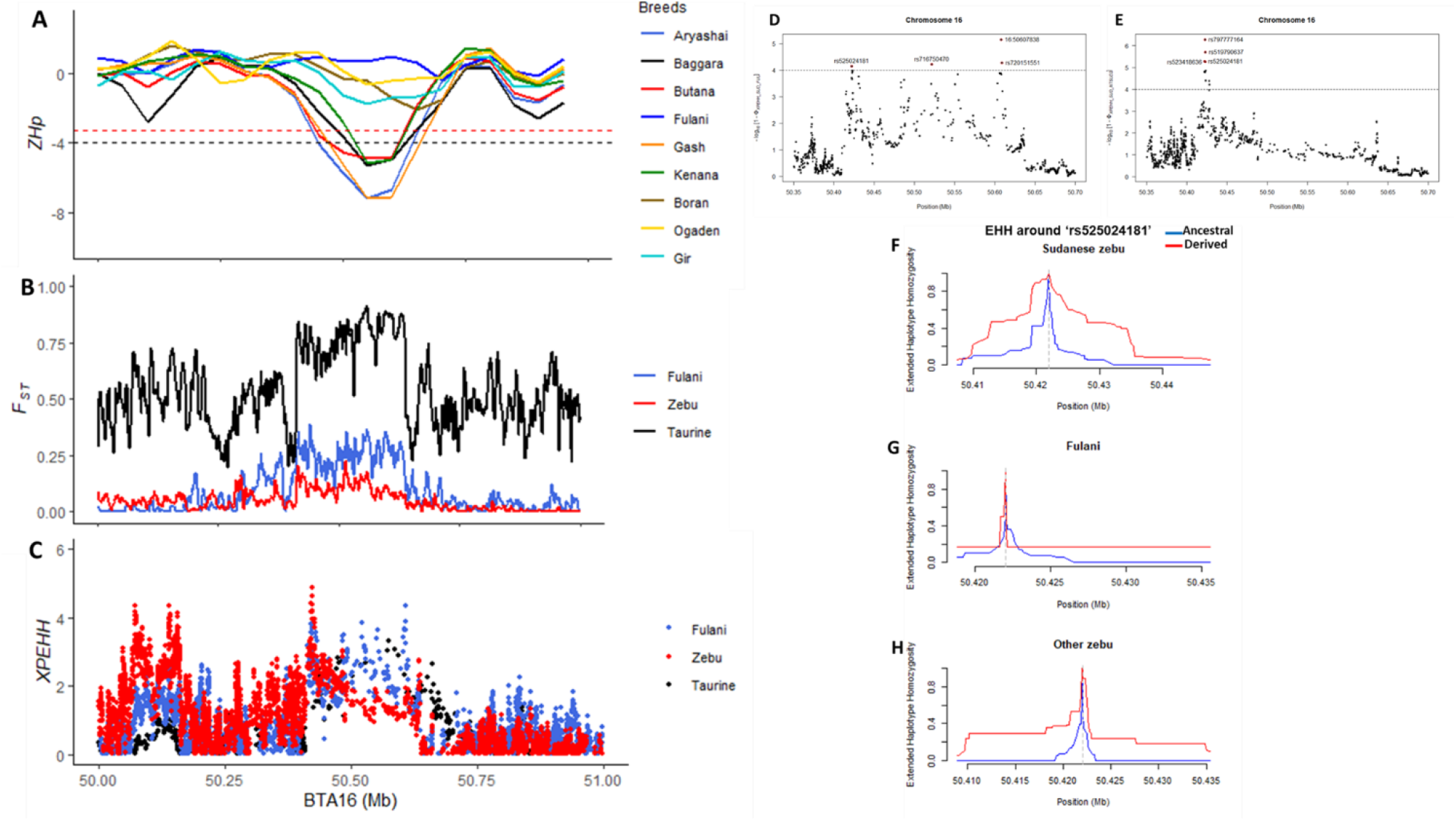
Strong evidence of positive selection in Sudanese zebu population at 250-kb chromosome 16 locus. **(A)** Genomic footprints of low diversity (*ZHp*) in the candidate region in eight African zebu breeds and one Asian zebu breed. The red and black dashed line indicates the maximum and minimum genome-wide *ZHp* selection outlier threshold values among the cattle breeds. **(B)** Population divergence (*F*_ST_) between the Sudanese zebu populations (excluding the Fulani) and other zebu and taurine cattle populations. **(C)**Compared to other cattle populations, there was increased haplotype homozygosity (XP-EHH) at a 250 kb selective sweep locus in Sudanese zebu. **(D, E)** Localisation of SNPs at the selective sweep region following XP-EHH between the combined five Sudanese breeds and Fulani **(D)**, as well as other zebu breeds (East African and Asian breeds) **(E)**. **(F, G, H).**The decay of extended haplotype homozygosity around rs525024181 in Sudanese Zebu **(F)**, Fulani **(G)** and other zebu breeds (**H**).

### A unique Sudanese zebu-specific candidate selective sweep at chromosome 16 locus

To further confirm the evidence of recent selection at the chromosome 16 locus and possibly avoid the problem of confounding demographic factors (Oleksyk *et al*., 2010), we performed additional genomic scan analysis on the entire bovine chromosome 16 using the XP-EHH test (Sabeti *et al*., 2007). We used the Sudanese Fulani, the other zebu breeds and the taurine population as three separate control groups (Figure S2A-C). The haplotype homozygosity in the Sudanese zebu population on the chromosome 16 locus is shown in Figure 6C. In particular, the comparison with Fulani reveals selection signals at four significant SNPs (-log10 *XP-EHH*, > 4), which are located within three genes (*PRKCZ* (16:50607838 and rs720151551), *SKI* (rs716750470) and *PLCH2* (rs525024181) (Figure 6D). In contrast, XP-EHH comparison with other zebu populations (East African and Asian breeds) reveals one strong signal containing several significant SNPs (Figure 6E). The latter’s top four significant SNPs (rs797777164, rs519790637, rs525024181 and rs523418636) are located within the *PLCH2* gene.

Consequently, *PRKCZ*, *SKI* and *PLCH2* genes might be zebu-specific candidate genes linked to local adaptation to the dryland habitat in Sudan’s desert and semi-desertic regions. Indeed, they are involved in alternative metabolic processes of glucose homeostasis, insulin signalling and fat metabolism, which are likely relevant adaptive strategies in hot and arid environments (Diaz *et al*., 2012, Zou *et al*., 2013, Lagundžin *et al*., 2019, Ahmed *et al*., 2020, Rocha *et al*., 2021). Furthermore, we considered the one common significant SNP (rs525024181, G>A) within the *PLCH2* genes (Figure 6D-E) as a putative candidate variant or closely linked polymorphism targeted by selection. The haplotype decay around rs525024181 reveals that the haplotype containing the derived allele has more extensive homozygosity in the Sudanese zebu than in the other zebu breeds (Figure 6F - H), suggesting a strong selection at the locus in the Sudanese population. We did not observe much difference in the pattern of EHH around the remaining seven significant SNPs among the zebu populations (Figure S3A – C).

## Discussions

In this study, we analyse for the first time the whole-genome sequences of six indigenous African zebu breeds from the most extreme African dryland conditions of Sudan. We aimed to identify candidate selective sweep regions linked to dryland adaptive traits and to assess their uniqueness among other African and non-African cattle. We adopted three genomic scan approaches to detect genomic regions of reduced diversity (*ZHp*), population divergence (*F*_ST_), and increased haplotype homozygosity (XP-EHH). Most of the within-breed *ZHp* genomic signatures detected are unique to individual zebu breeds. Interestingly, we find several gene-rich regions among the shared candidate selective sweeps across Sudanese or between Sudanese and other zebu breeds. In addition, some of these regions also show high population divergence signals between the Sudanese zebu and other zebu and taurine breeds. These results are compatible with the identified selective sweeps being *Bos indicus* specific, while others being rather African dryland specifics as detected exclusively in the Sudanese cattle population of the arid environments. If under common regulatory control, these gene-rich regions will be consistent with the view that change in gene expression may control complex traits (Naval-Sanchez *et al*., 2018) while providing a possible mechanism for rapid adaptation to new environmental challenges (Gheyas et al. 2021)

Interestingly, the functions of the genes within candidate gene-rich regions identified in the present study are associated with phenotypes such as body size and conformation, stress response to heat, immune response, insulin signalling, glucose metabolism, and fat metabolism. Interestingly, positively selected genes involved in similar processes have been reported in other desert-dwelling mammals like sheep, goats and camels. However, many reported genes are not the same (Wu *et al*., 2014, Kim *et al*., 2016, Mwacharo et al., 2017). Nevertheless, the results here support the evidence of common adaptive traits or mechanisms among the ungulates adapted to hot and arid climates (Mirkena *et al*., 2010, Gebreyohanes *et al*., 2017, Rocha *et al*., 2021).

Among the highly frequent *Bos indicus*-specific, selective sweeps are candidate genes involved in body size, conformation, stress response to heat and immune response. Notably, the *HMGA2* gene within the expanded selective sweep on chromosome 5 has been linked to different phenotypes, including growth and conformation in other non-African cattle breeds (Porto-Neto *et al*., 2014, Yurchenko *et al*. 2018, Aguiar *et al*., 2018, Naval-Sánchez *et al*., 2020, Dixit *et al*., 2021), and other species (Kim *et al*., 2004, Weedon *et al*., 2007, Boyko *et al*., 2010, Fariello *et al*., 2014 Sevane *et al*., 2017 and Dutta *et al*., 2020). Interestingly, for the first time, we identified strong selection signatures in the region of this pleiotropic gene in the indigenous African cattle population. On the contrary, several genes overlapping the extended gene-rich selective sweep on chromosome 7 have previously been detected in African and non-African zebu breeds (Kim *et al*., 2017, Bahbahani *et al*., 2017, Kim *et al*., 2020, Naval-Sánchez *et al*., 2020. In particular, Kim *et al*. (2020) reported an excess of indicine ancestry in this region. Furthermore, among the twenty-eight genes in this region is the DnaJ heat shock protein family (HSP40) member C18 (*DNAJC18*), also reported in tropically adapted sheep breeds from Ethiopia (Ahbara *et al*., 2019) and Bactrian camel (Wu *et al*., 2014). Undoubtedly, the genes in this region are involved in stress response to heat. However, the region requires further investigation and fine mapping to identify the probable candidate gene(s) or variant(s) targeted by selection.

Exposure to infectious and parasitic diseases challenges is another stressor faced by Sudanese cattle living in hot arid environments, necessitating an adequate immune response for survival in the region. We found the interleukin 1 receptor-associated kinase 3 (*IRAK3*) gene in the proximity of the *HMGA2* gene. *IRAK3* is a crucial modulator of inflammatory responses and negatively regulates Toll-like receptor signalling pathways involved in innate host defence (Nguyen *et al*., 2020). It has been reported under positive selection in non-African humped cattle adapted to tropical climatic conditions (Porto-Neto *et al*., 2014; Naval-Sánchez *et al*., 2020). Another detected innate immune-related gene is the macrophage receptor with collagenous structure (*MARCO*), shared by most Sudanese breeds and the Gir cattle but not the African non-Sudanese zebu breeds. The gene is likely involved in immune responses against bacterial infection, particularly the causative agent of tuberculosis, *Mycobacterium bovis*. Specifically, *MARCO* plays a role in the phagocytosis of bacteria, including Mycobacterium species and Salmonella species, as demonstrated in previous studies involving microbial infections in mice and zebrafish (Dorrington *et al*., 2013; Benard *et al*., 2014). Polymorphisms in the gene have also been associated with pulmonary tuberculosis (TB) in humans caused by *Mycobacterium tuberculosis* (Ma *et al*., 2011; Bowdish *et al*., 2013; Thuong *et al*., 2016). Although *M. tuberculosis* is the causative agent of TB in humans, the incidence of this anthropogenic disease is frequently identified in cattle, particularly in Africa, including Sudan, where the disease’s prevalence is also high in humans. It is believed that the transmission between man and cattle results from the close association of cattle farmers to their animals (Ocepek *et al*., 2005; Thoen *et al*., 2008). Indeed, Sudan happens to be one of Africa’s regions where a very high prevalence of TB in cattle herds has been reported (Sulieman and Hamid 2002; Ibrahim *et al*., 2016). Also, the only two anecdotally known African dairy zebu breeds, Butana and Kenana, are indigenous to Sudan. This gene’s role in TB susceptibility in humans and its detection in this study in a candidate region under positive selection in African cattle further supports cattle as asymptomatic carriers of the human’s *Mycobacterium tuberculosis*.

In addition to extreme temperature and disease challenges, other harsh conditions of the African dryland areas include food and water shortages (Jonsson 2006, Uddin and Kebreab 2020). Indeed, we identified a unique Sudanese zebu selective sweep on chromosome 16 detected exclusively in five Sudanese breeds (except the Fulani). Furthermore, this region shows high population differentiation between the Sudanese zebu and other cattle populations, including the Sudanese Fulani. The absence of this selection signal in the Fulani will need to be further investigated, considering the genetic closeness of the Fulani and Baggara and the origin of the Fulani breed, predominantly found in the West African Sahelian region (Rahman 2007, Nahar 2009). The candidate genes overlapping this unique sweep have roles in glucose homeostasis, feed efficiency, insulin signalling and fat metabolism. Glucose homeostasis and insulin signalling are essentially biological processes contributing to animals’ survival due to nutritional shortages in the desert. Also, there are pieces of evidence of the periods of dietary restriction in cattle and other species coinciding with increased insulin sensitivity and glucose uptake (Cartee *et al*., 1994, Röpke *et al*., 1994, Sternbauer and Luthman 2002, Barretero-Hernandez *et al*., 2010). In addition, *FAAP20, PRKCZ*, and *SKI* genes have been implicated in insulin metabolism (Leong *et al*., 2010, Diaz *et al*., 2012, Fitzsimons *et al*., 2014, Kriebel *et al*., 2016, Lagundžin *et al*., 2019). In particular, *PRKCZ* functions by regulating the translocation of the glucose transporter 4 (*GLUT4*) to the cell surface for glucose uptake and has been implicated in insulin response to glucose uptake in fasted cattle. Thus, a greater expression of both *PRKCZ* and *GLUT4* has been observed in fasted animals, suggesting a greater sensitivity to glucose uptake by myocytes (Keogh *et al*., 2015).

Interestingly, consistent with previous reports on the profound increased expression levels of *GLUT1* (glucose transporter 1) and genes involved in glycolysis in the renal medulla of water-deprived Bactrian camels (Wu *et al*., 2014). Thus, our results suggest shared physiological responses facilitated by glucose transporter proteins in camelids and cattle during water and food scarcity. However, the processes may have evolved more rapidly in camelids than in cattle as a result of *GLUT1* being ubiquitously expressed and possessing high capacity than the *GLUT4*, which is expressed mainly in insulin-sensitive tissues like the heart, skeletal muscle and adipose tissue (Scheepers *et al*., 2004, Wu *et al*., 2014). In addition, increased expression of the insulin-responsive glucose transport proteins has been correlated with improved insulin action in skeletal muscle, especially during exercise. Indeed, overexpression of the *SKI* gene in the skeletal muscle has been demonstrated to modulate the genetic controls of insulin signalling and, thus, glucose homeostasis (Stallknecht *et al*., 2000, Diaz *et al*., 2012).

The above function may correlate with whole-body fat reduction in exercised humans due to increased insulin sensitivity of triglyceride lipolysis in subcutaneous adipose (Stallknecht *et al*., 2000). Also, data from transgenic animals have further supported the importance of adipose tissue in modulating whole-body insulin sensitivity (Shepherd *et al*., 1993). Hence, the role of adipose tissue mediated by lipid metabolism in the physiological adaptation process to hot arid environments has been well documented. Furthermore, adipose tissue serves as an organ for storing food and a source of energy and water, contributing to animal survival during prolonged starvation and thirst. Moreso, fat oxidation produces metabolic water in water-deprived or exercised animals (Pond 2017). Strikingly, two other genes, *PLCH2* and *PEX10*, found within the unique Sudanese zebu selective sweep on chromosome 16, are involved in the adaptive metabolic strategy of lipid catabolism and oxidation. Functionally, the phospholipase C eta 2 (*PLCH2*) gene, also known as *PLCL4*, among other aliases, is suggested to participate in the lipid catabolic process for glucogenesis. Moreover, the hypermethylation of loci associated with *PLCH2* coinciding with fat reduction has also been reported in calory restricted humans (Bouchard *et al*., 2010). On the other hand, peroxisomes such as *PEX10* controls the composition of intracellular fatty acid content especially, the unsaturated fatty acid content, and are essential in the oxidation of fatty acids to produce water (Hong *et al*., 2009, Fillmore *et al*., 2011, Yan *et al*., 2020).

In conclusion, the present study analyses the whole genome re-sequencing of six indigenous Sudanese zebu breeds to understand their genome diversity and the genomic footprints of their adaptations to environmental challenges in the African dryland area. We reported several gene-rich selective sweeps following three genomic scan approaches. The results highlight the importance of selection at gene-rich genome regions as a mechanism of adaptation to the complexity of environmental challenges. If under the genetic control of a similar regulatory mechanism, these gene-rich regions may contribute to the rapid adaptation of cattle to new challenging environments. Also, these regions could be enriched for superalleles, but this will require further investigation. Moreover, we identified a unique selective sweep on chromosome 16 overlapping multiple genes involved in insulin signalling, feed efficiency, glucose and fat metabolism, strongly suggesting their roles in adaptive metabolic strategies during feed and water deprivation and energy requirements. Our results align with previous studies that reported the relevance of these cellular processes in other species, including humans, enabling adaptation to similar conditions of the harsh desert environment.

## Supporting information

https://drive.google.com/drive/folders/1F67HF8F1MG-GiSrN4gvLyvXzjiplaAc5?usp=sharing

## Author contributions

Author contributions: HM, OH, and AT designed the research; HM secured funding for the sampling and sequencing of the Group A Sudanese zebu cattle; MVB contributed the data on Gir cattle from Brazil; BS, HE and TM collected the Group A Sudanese samples; KM and BS collected and contributed the Group B Sudanese zebu samples; AT analysed the data; AT and OH interpreted the results; AT drafted the manuscript, AT and OH edited the manuscript; All authors revised and approved the manuscript’s final version.

## Acknowledgements

The Group A Sudanese zebu breeds were preliminary analysed and presented in the University of Nottingham thesis of Dr Abdulfatai Tijjani. The authors acknowledge the Ministry of Finance and National Economic, Sudan, and the African Development Bank, country office (Khartoum, Sudan) to support the project’s approval and implementation.

## Supplementary Figures

**Figure S1.**
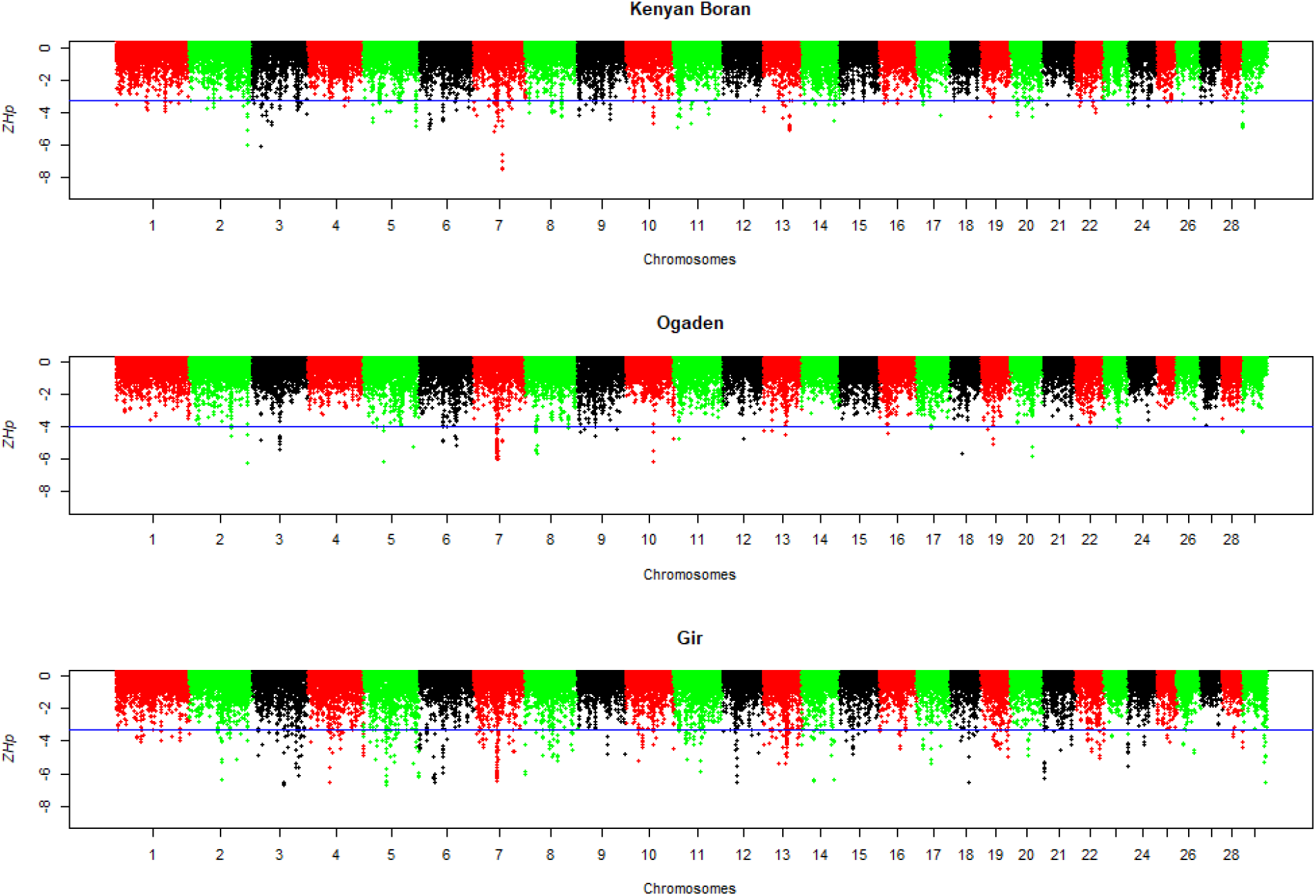
The distributions of Z-transformed Hp scores across the bovine autosome in three zebu breeds (non-Sudanese); Kenya Boran (A), Ethiopian Ogaden (B) and Gir from Brazil (C). The blue line indicates the *ZH*p threshold value (lowest 0.5%) for selecting outlier windows (candidate regions under positive selection).

**Figure S2.**
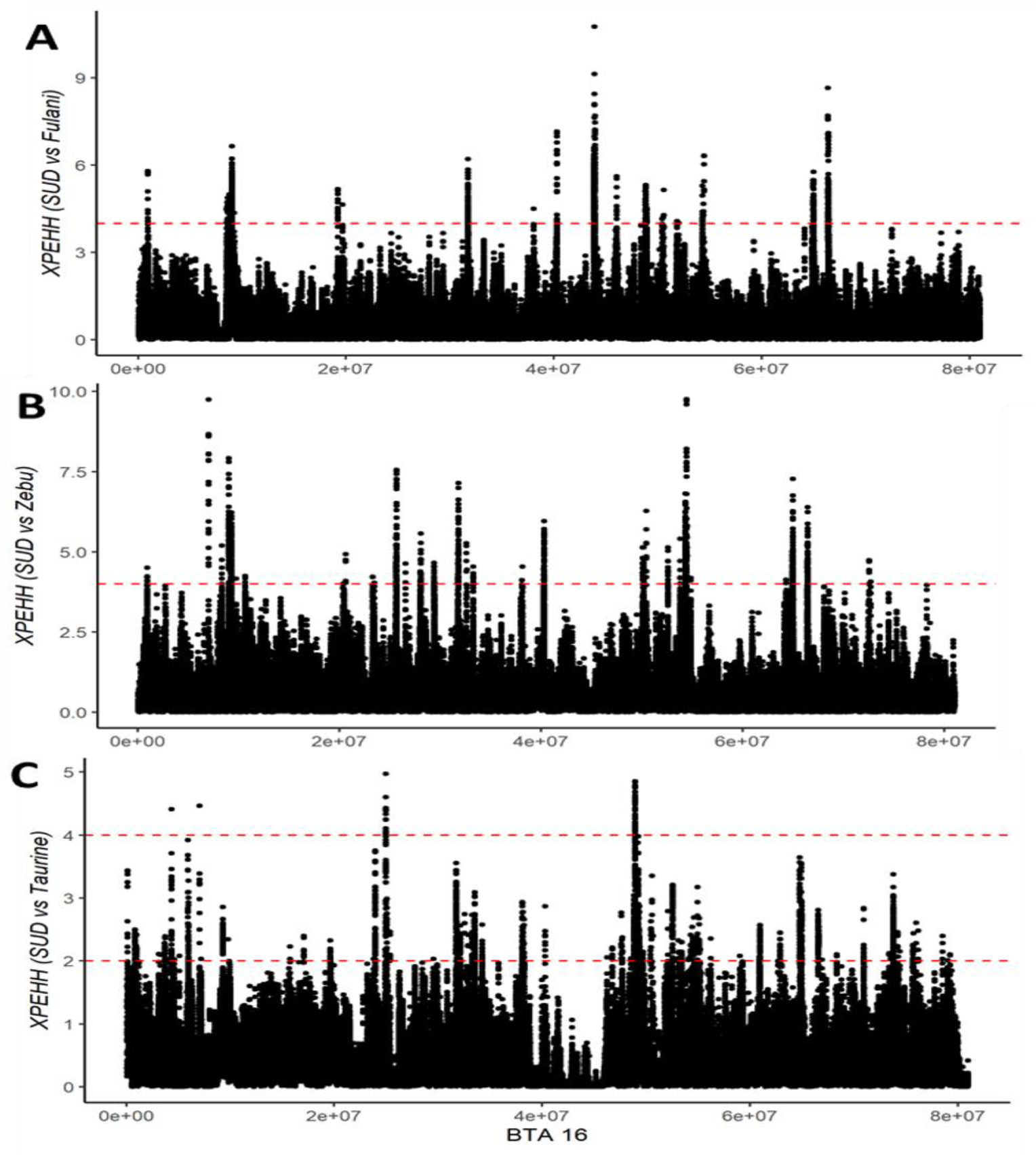
The distribution of XP-EHH scores along Bovine chromosome 16 following the comparison of Sudanese zebu population with other zebu and taurine cattle populations. Sudanese Fulani (A), other zebu breeds (East African and Asian zebu) (B), and Taurine breeds (West African and Euroasia and West Europe breeds) (C). The red dash blue line indicates the significant XP-EHH threshold of >4 (*p*_value <0001) for selecting outlier SNPs (SNPs targeted by positive selection).

**Figure S3A.**
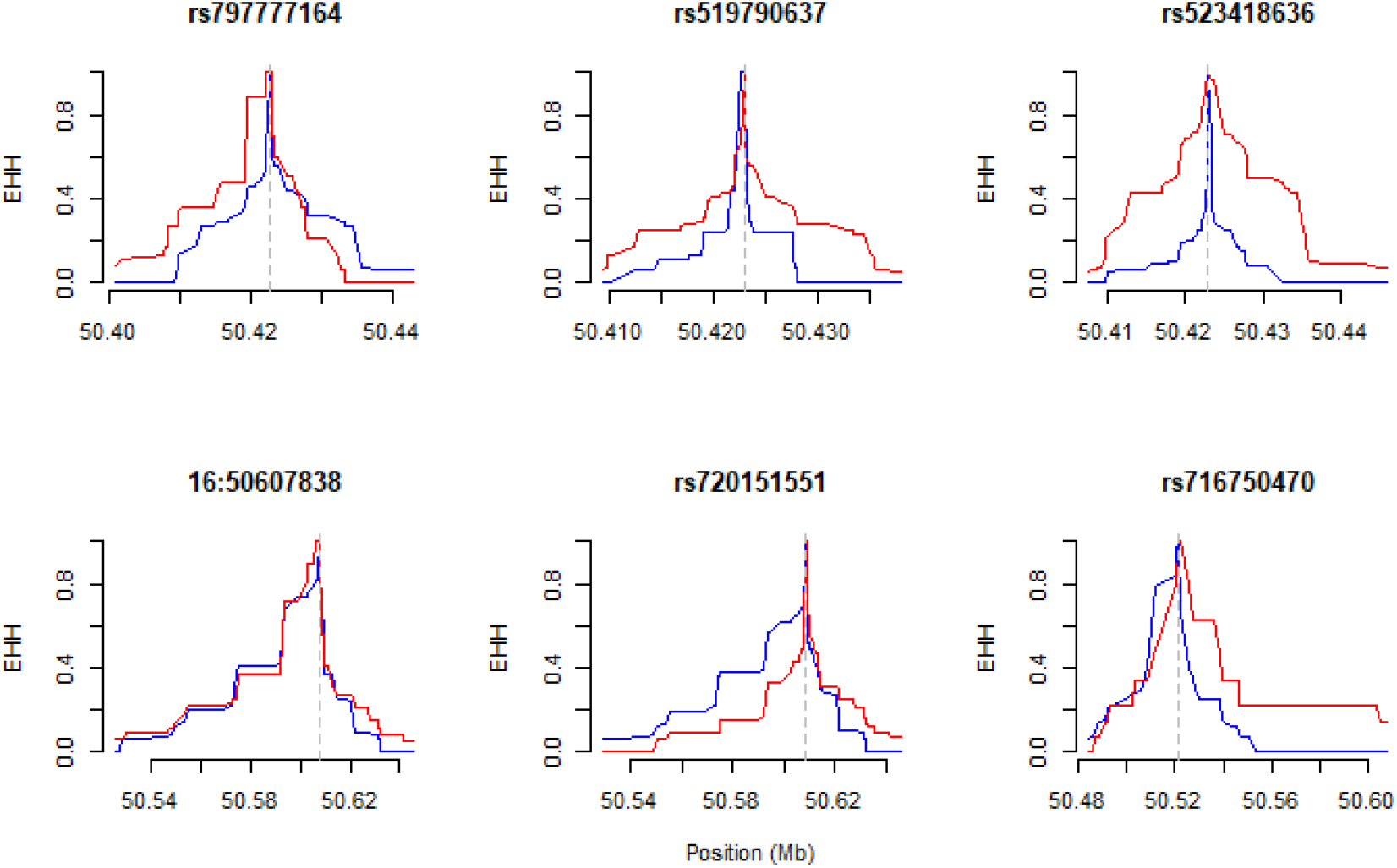
The decay of extended haplotype homozygosity around six significant SNPs in the Sudanese zebu population

**Figure S3B.**
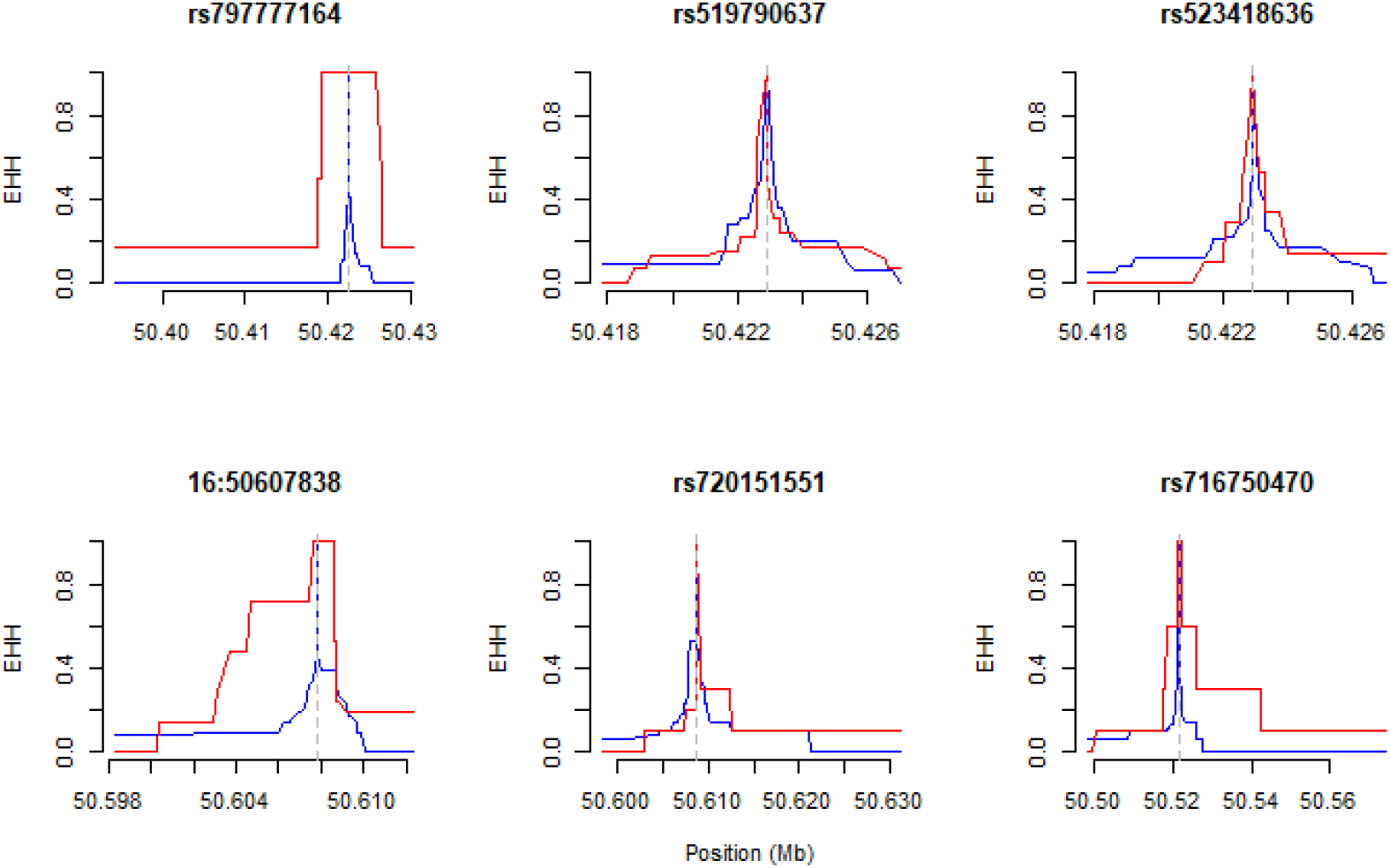
The decay of extended haplotype homozygosity around six significant SNPs in Sudanese Fulani

**Figure S3C.**
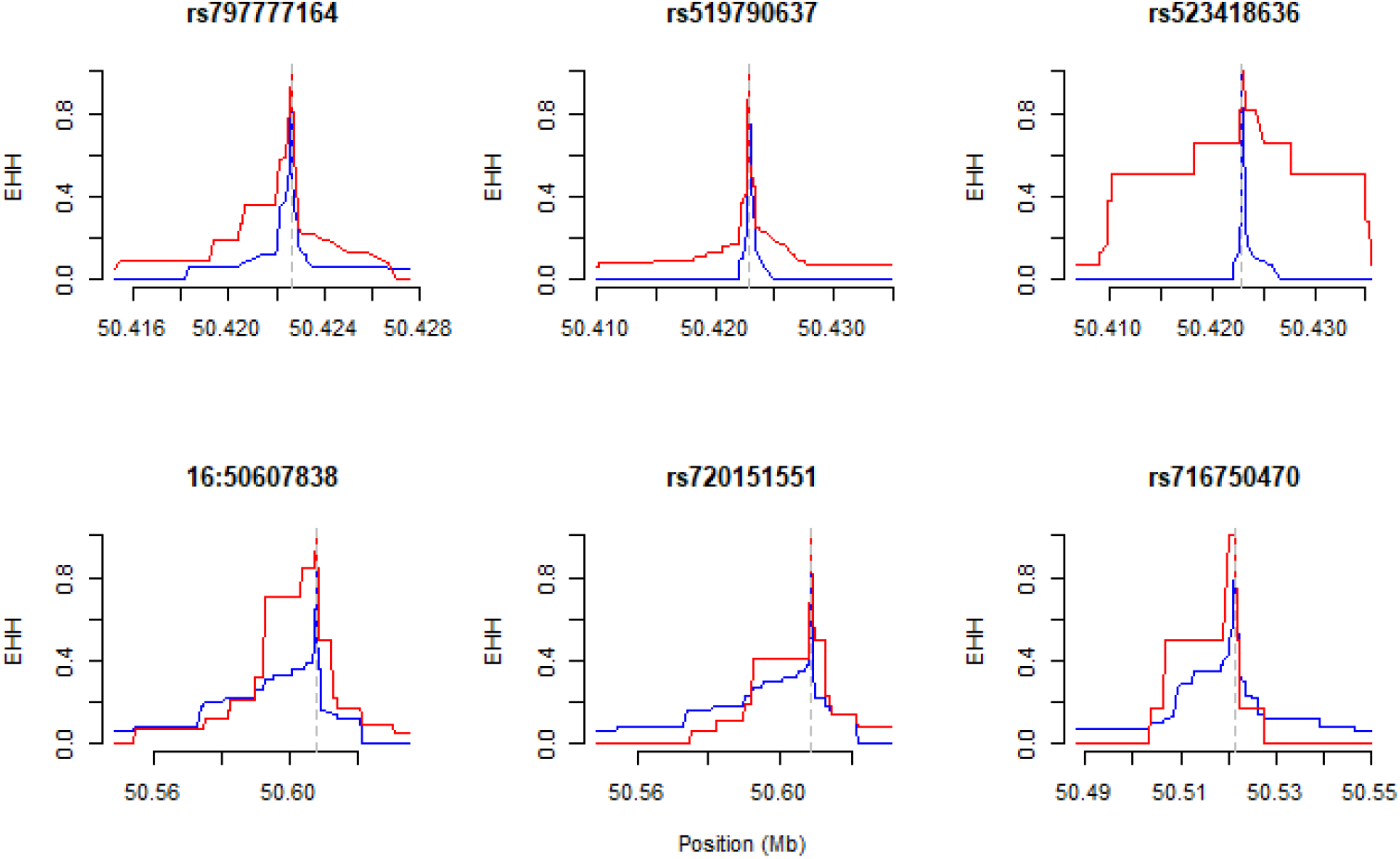
The decay of extended haplotype homozygosity around six significant SNPs in other zebu population

